# Semi-Self-Supervised Learning for Semantic Segmentation in Images with Dense Patterns

**DOI:** 10.1101/2022.08.09.503251

**Authors:** Keyhan Najafian, Alireza Ghanbari, Mahdi Sabet Kish, Mark Eramian, Gholam Hassan Shirdel, Ian Stavness, Lingling Jin, Farhad Maleki

**Affiliations:** Department of Computer Science, University of Saskatchewan, Saskatoon, Saskatchewan, Canada; Mathematics Department, Faculty of Sciences, University of Qom, Qom, Iran; Department of Mathematics, Faculty of Mathematical Science, Shahid Beheshti University, Tehran, Iran; Department of Computer Science, University of Calgary, Calgary, Alberta, Canada

## Abstract

Deep learning has shown potential in domains where large-scale annotated datasets are available. However, manual annotation is expensive, time-consuming, and tedious. Pixel-level annotations are particularly costly for semantic segmentation in images with dense irregular patterns of object instances, such as in plant images. In this work, we propose a method for developing high-performing deep learning models for semantic segmentation of wheat heads utilizing little manual annotation. We simulate a computationally-annotated dataset using a few annotated images, a short unannotated video clip of a wheat field, and several video clips from fields with no wheat. This dataset is then used to train a customized U-Net model for wheat head segmentation. Considering the distribution shift between the simulated and real data, we apply three domain adaptation steps to gradually bridge the domain gap. Only using two annotated images, we achieved a Dice score of 0.89 on the internal test set, i.e., images extracted from the wheat field video. The model trained using only two annotated images was evaluated on a diverse external dataset collected from 18 different domains across five countries and achieved a Dice score of 0.73. To further expose the model to images from different growth stages and environmental conditions, we incorporated two annotated images from each of the 18 domains and further fine-tuned the model. This resulted in improving the Dice score to 0.91. These promising results highlight the utility of the proposed approach in the absence of large-annotated datasets. Although the utility of the proposed method is shown on a wheat head dataset, it can be extended to other segmentation tasks with similar characteristics of irregularly repeating patterns of object instances.

## 1 Introduction

Deep learning models have shown promising results in various computer vision tasks, including object recognition [1], object detection [2], instance segmentation [3], and semantic segmentation [4]. Deep learning has shown the potential to be widely utilized in precision agriculture, which focuses on computational methods to sustainably improve the quality and quantity of crop production [5–12]. The availability of a large-scale and diverse dataset like the Global Wheat Head Detection (GWHD) dataset [13] has enabled researchers to develop novel supervised deep learning-based methods for detecting and counting wheat heads from field images [11, 14, 15]. However, the assessment of many important plant traits, such as organ size, organ health, biotic and abiotic stress, requires fine-grain semantic segmentation of plant organs. Semantic segmentation in plant images remains a significant challenge because creating pixel-level annotations for plant images is costly, due to the dense, partly occluding, and repeating pattern of plants and plant organs in field images.

For object detection in plant images, several recent studies have focused on developing fully supervised deep learning approaches for wheat head detection using the GWHD dataset [13]. Gong et al.[16] customised a YOLO [17, 18] model by introducing a dual spatial pyramid polling [19] and CSPNet [20] to enhance the detection speed and accuracy. Liu et al. [21] studied the importance of a linear dynamic color transformer in developing deep convolutional models for wheat head detection and reported that it could alleviate false negative predictions. Khaki et al. [14], utilized point-level annotation to develop a computationally lightweight deep model for wheat head detection based on a truncated version of MobileNetV2 [22].

In addition to object detection in agricultural applications, plant and leaf segmentation using deep convolutional neural networks has also gained considerable attention for feature extraction and pixel-level classification. Rawat et al. [23] studied the utility of active learning approaches—including pool-based active learning and four methods based on least confidence, margin, entropy, and deep Bayesian—for plant organ segmentation. Evaluating these approaches using three datasets for apple, rice, and wheat, they reported that random sampling outperformed active learning in two datasets. For wheat head segmentation, they manually segmented images from the UTokyo_Wheat_2020 dataset, which is a subset of the GWHD dataset containing high-quality images captured under good lighting conditions. Their model achieved an IoU of about 0.70 when evaluated on the data from the the UTokyo_Wheat_2020 dataset.

Hussein et al. [24] developed a deep learning-based segmentation pipeline for leaf segmentation in herbarium specimen images. They used DeepLabv3+ [25] with a pre-trained ResNet101 backbone [8]. Utilizing the connected components algorithm, they split the contour masks into individual leaves. Focusing on the segmentation of individual intact herbarium leaves for trait extraction, they used a single-leaf binary classifier based on a VGG16 network [26]. Their work showed that deep learning could be used for leaf segmentation and trait data extraction from herbarium specimen images.

Alkhudaydi et al. [27] utilized a Fully Convolutional Network [28] for wheat spike region segmentation. Using a small dataset of 90 side view images, they achieved an intersection over union (IoU) of 0.40 for wheat spike segmentation. The challenge with side view images is that the wheat heads in the middle of a plot often are not visible, making the utility of such a model very limited.

In the absence of large-scale annotated datasets for wheat head segmentation, several works have converted the wheat head segmentation or counting problem to a superpixel classification problem [29–32]. The main idea behind these works is to group pixels into homogeneous superpixels using the simple linear iterative clustering [33]. These superpixels then are classified to determine regions belonging to wheat heads. The superpixel generation is often sensitive to background clutter and illumination condition. Consequently, some superpixels partially cover spikes and non-spikes regions. This results in coarse segmentation masks.

Despite the success of supervised semantic segmentation models, the lack of availability of annotated datasets is still an obstacle to developing high-performing deep learning models for many domains. Therefore, methodologies that benefit from advances in supervised deep learning and also require a smaller amount of data annotation are of great value and could substantially expand the utility of deep learning models.

Unsupervised, self-supervised, and semi-supervised learning methods have been developed to alleviate the need for large-scale annotated datasets [34]. In unsupervised learning, the goal is to learn a compact data representation using unannotated data. Semi-supervised learning [35] aims at utilizing both annotated and unannotated data for model development. Self-supervised learning refers to techniques that, instead of relying on manual annotation, utilize supervisory signals that are computationally generated from the data [36].

In self-supervised learning methods, a pretext task—which is often different from the primary task—is first defined. Image rotation [37], Image inpainting [38], and Jigsaw puzzle [39–41] are examples of pretext tasks. The pretext task is designed to generate a computationally-annotated dataset. Then a supervised approach is used to develop a model using the computationally-annotated dataset. The rationale behind self-supervised learning is that a model while learning the pretext task learns low-level features that can be shared across tasks. This model can then be fine-tuned to learn the primary task using a smaller amount of annotated data.

Self-supervised and semi-supervised learning approaches have also been used for wheat head detection. Fourati et al. [42] developed a wheat head detection method based on Faster R-CNN [9], and EfficientDet [43] models. They used the GWHD dataset as the training dataset. They also utilized pseudo-labeling, which is considered a semi-supervised approach, alongside test time augmentation, multi-scale ensemble, and bootstrap aggregating to further increase the model performance, achieving a mean average precision of 0.74. Najafian et al. [44] proposed a deep learning-based approach, utilizing semi-supervised and self-supervised concepts, for wheat head detection. Developing a simulated dataset, they trained a YOLO architecture for wheat head detection using the simulated dataset. Their final model trained on the GWHD dataset achieved a mean average precision of 0.82, where a predicted bounding box with more than 50% overlap with the ground truth was considered an accurate detection.

Most segmentation tasks in precision agriculture share characteristics that makes them different from general segmentation tasks such as those developed using Pascal VOC [45] and MS COCO [46] datasets. Unlike general object segmentation, in precision agriculture most often, we are interested in segmenting densely packed and overlapping instances that are highly self-similar. In these applications, images often contain many repeated irregular patterns such as plant spikes, fruits, flowers, or leaves. Figure 1 illustrates a few examples of such images. These differences pose challenges and opportunities that should be considered when developing segmentation models in agricultural domains.

**Figure 1:**
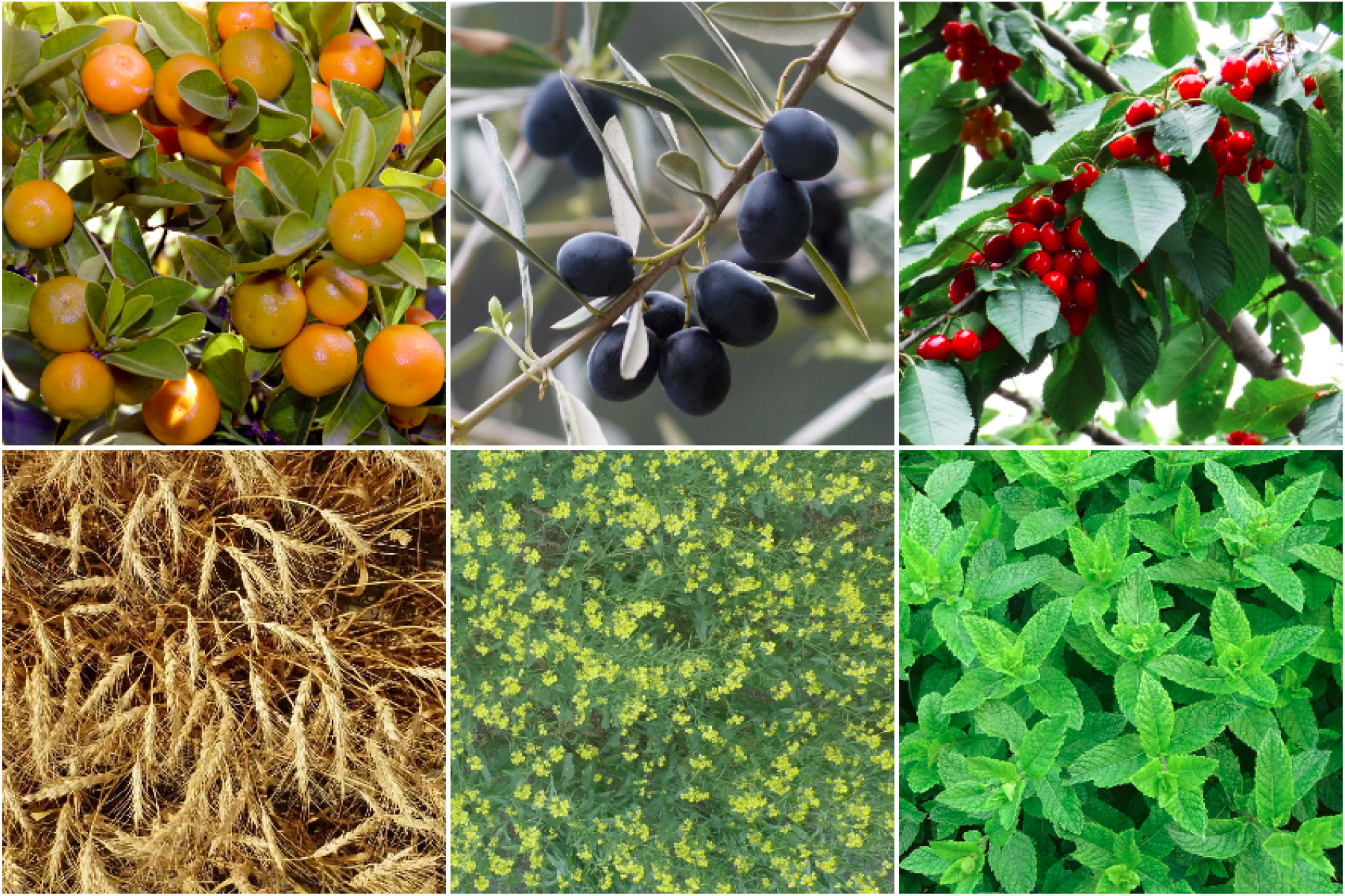
In agricultural images, we are often interested in densely packed, overlapping, and highly self-similar instances. In these applications, images often contain irregular patterns such as plant spikes, fruits, flowers, or leaves.

Considering these characteristics, in this paper, we develop a semi-self-supervised method that utilizes a short video clip of a wheat field, a few annotated images, and several short video clips of background scenes—i.e., fields with no wheat—to develop a wheat head semantic segmentation model. The proposed method utilizes a few annotated images to simulate a large-scale computationally-annotated dataset. Then, using this dataset, a customized U-Net model for wheat head segmentation is trained in a supervised manner. Finally, to address the domain gap between simulated and real images, three domain adaptation steps are applied. We evaluate the proposed method using the GWHD dataset, which is a diverse collection of wheat field images. The proposed method is a self-supervised approach because it relies on developing computationally-annotated data for model development. It is also a semi-supervised learning approach because it utilizes a few manually annotated images as well as pseudo-labeling for model training. The proposed approach alleviates the need for creating a costly, large-scale annotated dataset and facilitates the accelerated development of deep learning models for precision agriculture.

## 2 Materials and Methods

In this section, we present the methodology for developing wheat head segmentation models utilizing a small amount of annotated images. We first simulate a large-scale computationally-annotated dataset using a few annotated images. The synthesized images are then used to train a customized U-Net model for wheat head segmentation. Due to the difference between the synthesized and real images, also known as distribution shift, we expect the performance of the model trained on synthetic images to degrade when applied to real images. To address this domain gap, we apply three domain adaptation steps. We evaluate the proposed method using a diverse collection of wheat field images. In the following, we provide a detailed description of each step of the proposed method.

### 2.1 Data

We utilize a short video clip of a wheat field and 11 short video clips of background scenes to simulate annotated images. These video clips were obtained using Samsung cameras with 12 and 48 Megapixels resolution. Figure 2 illustrates snapshots of these video clips. We also perform an external evaluation using the Global Wheat Head Detection (GWHD) dataset [13] which includes images of wheat fields from five countries and 18 different domains from various stages of growth. Since there is no annotation for the GWHD dataset, we randomly select 10% of images (365 samples) from the GWHD dataset and manually segment them for model evaluation. Figure 3 illustrates examples of the images from the GWHD dataset with their segmentation masks overlaid. From the remaining images in the GWHD dataset, we randomly select 36 images (2 images from each of the 18 domains) and manually segment them. These images are then used for the final step of domain adaptation.

**Figure 2:**
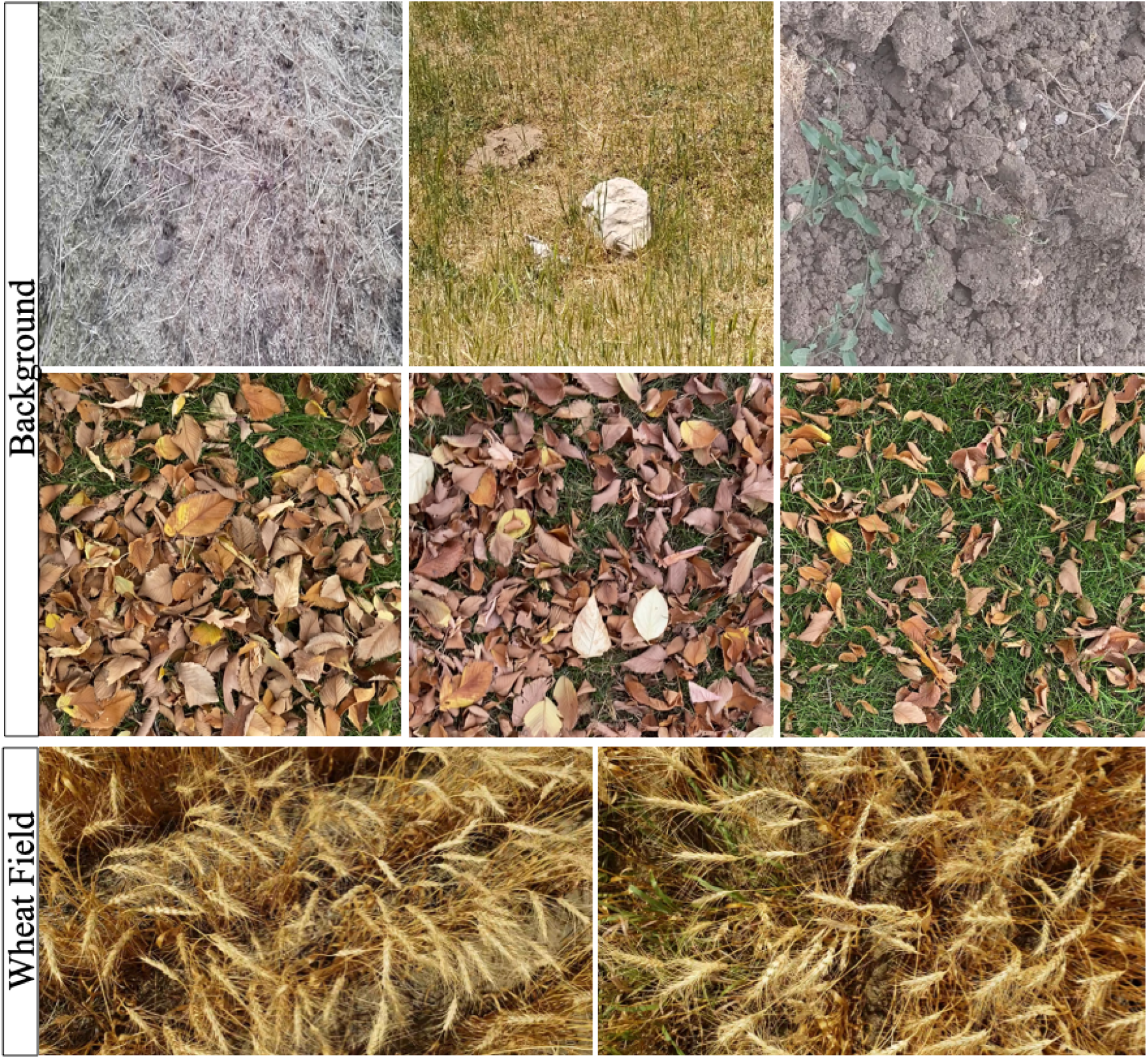
The top two rows show extracted image frames from the background video clips—i.e., video clips of fields with no wheat head. The third row shows examples of image frames extracted from the video clip of a wheat field.

**Figure 3:**
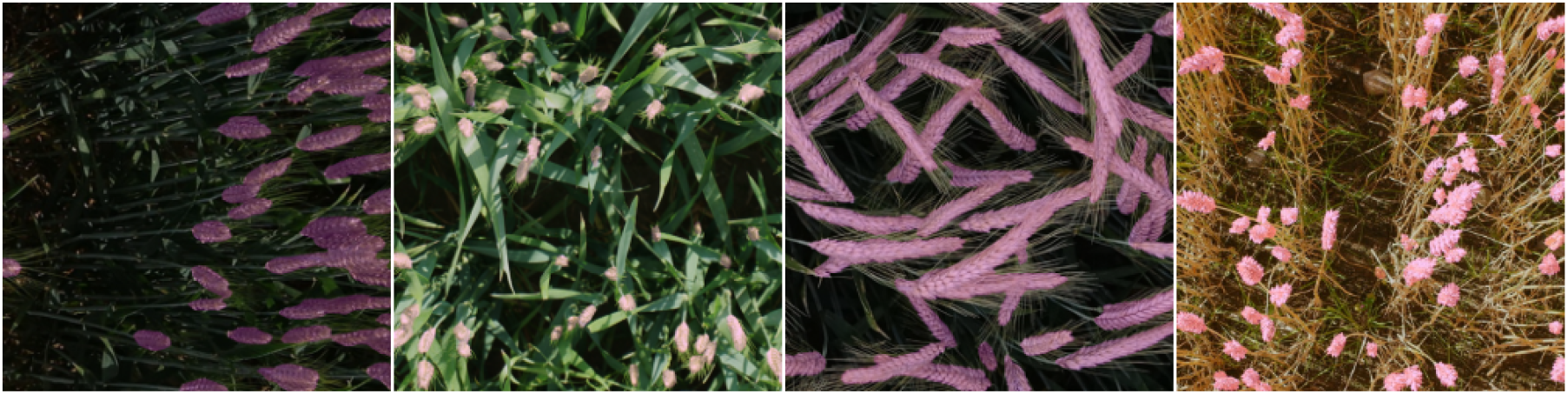
Examples of the GWHD dataset images manually annotated as the external test set. The segmentation masks are overlaid on the images.

### 2.2 Model

In this study, we use a customized U-Net [47] model architecture with the EfficientNet B4 [48] encoder pretrained on the ImageNet dataset [49, 50]. For all experiments in this paper, we employ the following loss function in Equation 1, which is the summation of binary cross-entropy and Dice loss functions.

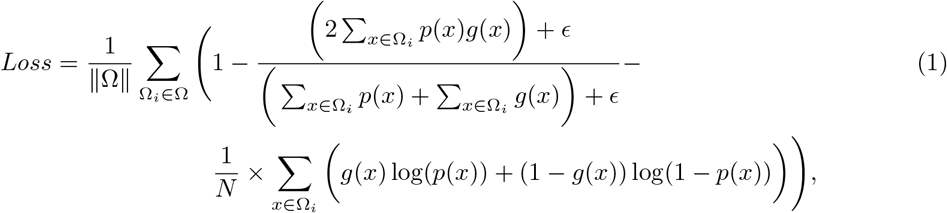

where *p*(*x*) is the probability of the pixel *x* being part of a wheat head on an image. *p*(*x*) is calculated by applying the sigmoid function to the output of the final convolution layer of the model. The function *g*(*x*) ∈ {0, 1} represents the ground truth class for pixel *x*, that is 1 if *x* is part of a wheat head and 0 otherwise. Ω is a batch of images in the training set, and Ω*_i_* is an image in Ω undergoing a sequence of image transformations with computationally inferable labeling functions.Also, *ϵ* is a smoothing constant to prevent division by zero. We used *ϵ* = 1*E* – 05 for all experiments.

Assume that for a data point *x* in a domain *X*, there is a labeling function *ψ* that assigns a label *y* to x, that is, *ψ*(*x*) = *y*. Also, assume that 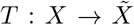 is a transformation function that maps each data point *x* from domain *X* to a data point 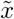 from domain 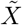. The labeling function *ψ* for a transformation *T* is computationally inferable if for any *x* ∈ *X*, there is a deterministic algorithm that outputs *ψ*(*T*(*x*)), which is the label for 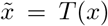. Note that we differentiate a sequence of image augmentations from a sequence of image transformations with computationally inferable labeling functions, as the former does not always lead to an image where labels can be inferred. For example, through the image augmentations, an image might get so distorted that even for an expert, manual labeling is not possible. We use the Albumentations package [51] for all image augmentations in this study.

In this study, we utilize the commonly used Dice score and intersection over union (IoU) as our performance measures [52, 53]. Given an observed segmentation mask *O* and the expected segmentation mask *E*, the Dice score is defined as:

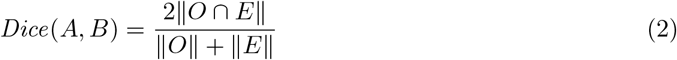

where ║ ║ and ∩ represent set cardinality and intersection operators, respectively. IoU is defined as follows:

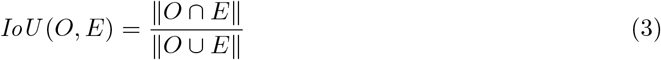

where ∪ represents the union operator.

In all experiments, we use the smallest loss value on the validation sets as the criterion for model selection. We also use the SGD optimizer [54] with a learning rate of 1e–2.

### 2.3 Data synthesization

We simulate a dataset of wheat field images using video clips of wheat fields and background scenes. We extract image frames from the video clips of the background fields to generate a set of *m* background images *B* = {*b*_1_, *b*_2_,…, *b_m_*}. We also extract image frames from the wheat field video clip to generate a set of *n* wheat field images *W* = {*w*_1_,…, *w_n_*}. We selected two different images, *w_t_* and *w_v_*, from the set *W* which are then used for simulating the training and validation subsets of the simulated dataset, respectively.

Figure 4 illustrates the process of simulating images using *w_t_, w_v_*, and background images. The procedure for simulating dataset *S_t_* is as follows. First, we manually segment the wheat heads in *w_t_*. This results in a set of wheat heads *H_t_*. We select a background image *b_i_* from *B*. This image serves as a canvas where we overlay: (1) a set of fake wheat heads, which only have the geometry of real wheat heads but are extracted from the background area of *w_t_* and (2) a set of real wheat heads from *H_t_* selected based on random sampling with replacement approach. The number of fake and real wheat heads to be randomly placed on *b_i_* is chosen as uniformly distributed random numbers between 10 and 100. We first overlay fake wheat heads and then real wheat heads. Also, prior to placement on *b_i_*, both real and fake wheat heads undergo a sequence of image augmentations including horizontal and vertical flips, rotation, resizing, and elastic transformation [55]. The locations of the real wheat heads are recorded to generate the segmentation mask for the simulated image. The resulting image undergoes a sequence of color augmentations [51].

**Figure 4:**
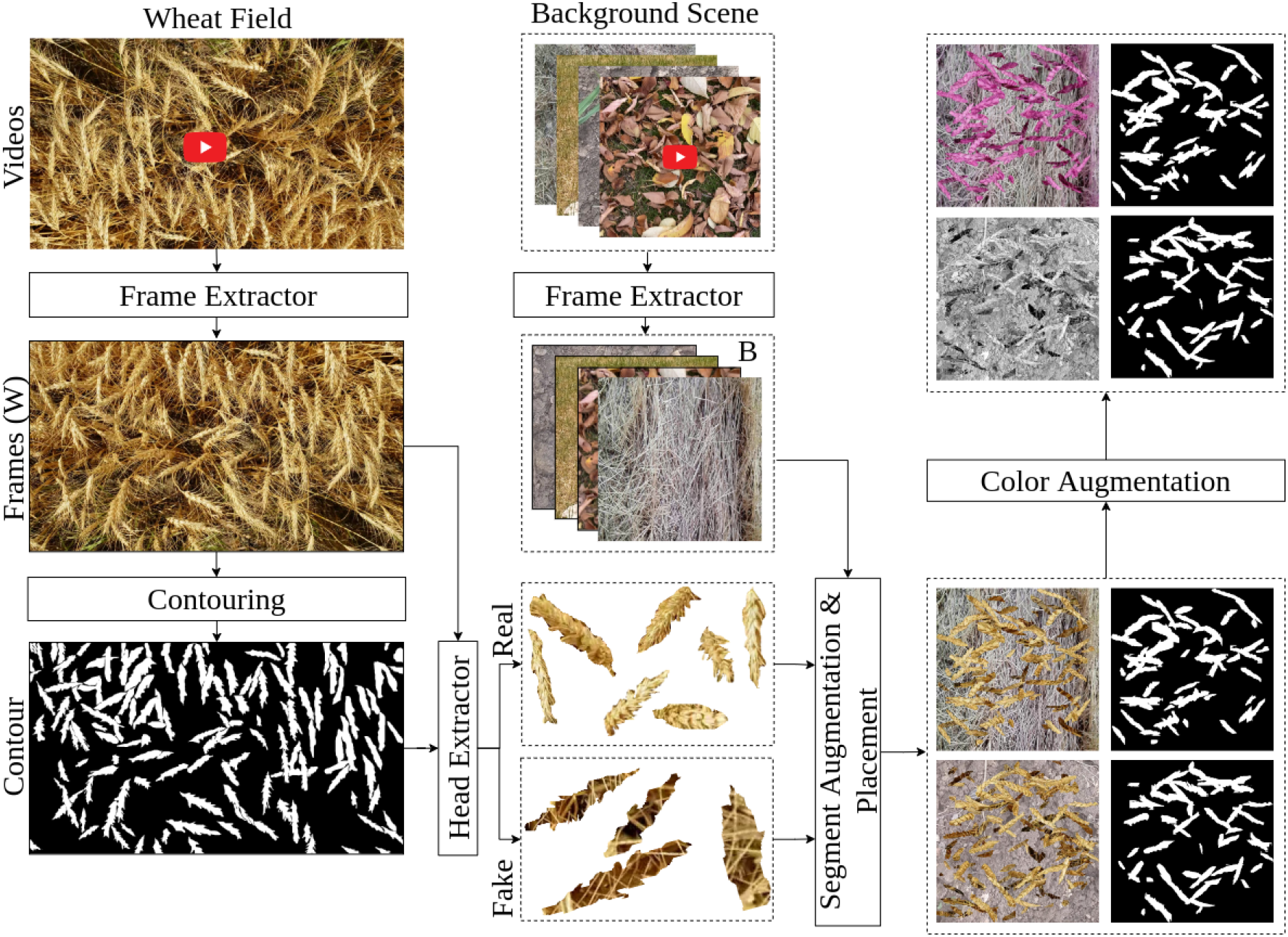
The procedure for simulating the computationally-annotated datasets—i.e., *S_t_* and S_v_.

We repeat this process 10,000 times to form *S_t_* with 10,000 simulated images. A simulated dataset *S_v_* of size 1000 is generated analogously. *S_t_* is used for model training, and *S_v_* is used for model evaluation. In the rest of the paper, a set with subscript *t* represents a training set, and a set with subscript *v* represents a validation set. We refer to the model trained with these datasets as model *S*. We train and evaluate the model for 10 epochs. Note that instead of developing a small dataset and using a larger number of epochs, we simulate a larger dataset and use a smaller number of epochs to avoid model overfitting. The rest of the models, which are developed for the domain adaptation purposes, are trained for 20 epochs (see below). Also, all models are trained using images of size 1024 × 1024 pixels.

We also use a test time augmentation (TTA) approach to improve the predictions made by the model. During this process, for each image, we generate two augmented versions using Gaussian noise and sepia image augmentations [51]. Then the predictions made by the model for the original image and its two augmented versions are aggregated using a pixel-level majority vote to compute the final prediction.

### 2.4 Domain adaptation

Since there is a domain shift between the images in the simulated datasets *S_t_* and *S_v_* and real images (as illustrated in Figure 6), a domain adaptation step is required to improve the model performance. In this paper, we apply three domain adaptation steps. As the first domain adaptation step, we develop datasets *D_t_* and *D_v_* that are semantically more similar to real images in comparison to the simulated images in *S_t_* and *S_v_* (see Figure 5) and use these datasets to fine-tune the model S. Figure 5 illustrates the process for generating images in datasets *D_t_* and D_v_. These datasets are generated by utilizing all 360 rotations (from 0 to 359 degrees) of *w_t_* and *w_v_*, respectively. These rotated images undergo a comprehensive set of image augmentations, which will be applied in an online manner—i.e., each time we access an image, a different sequence of image augmentations is applied to the image. These augmentations are applied to increase the data variability and avoid overfitting. Appendix A provides the list of image augmentations applied to images in *D_t_* and D_v_. We refer to the model trained with these datasets as model *D*.

**Figure 5:**
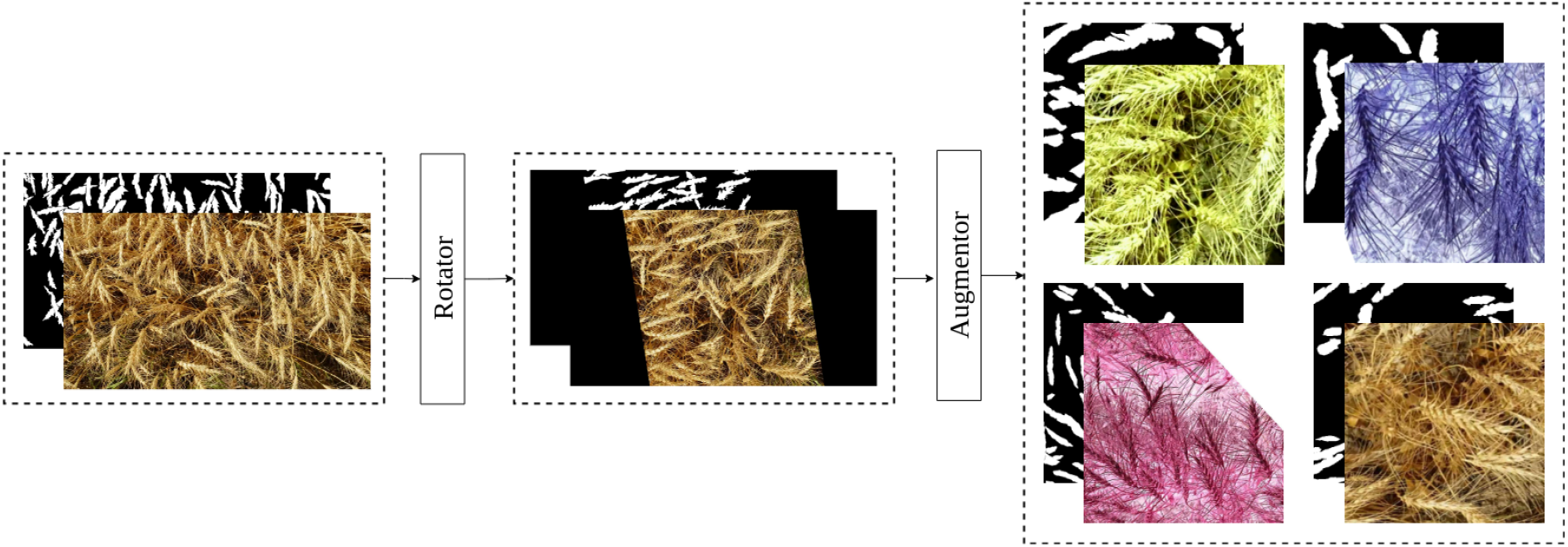
The images generated during the first domain adaptation step.

**Figure 6:**
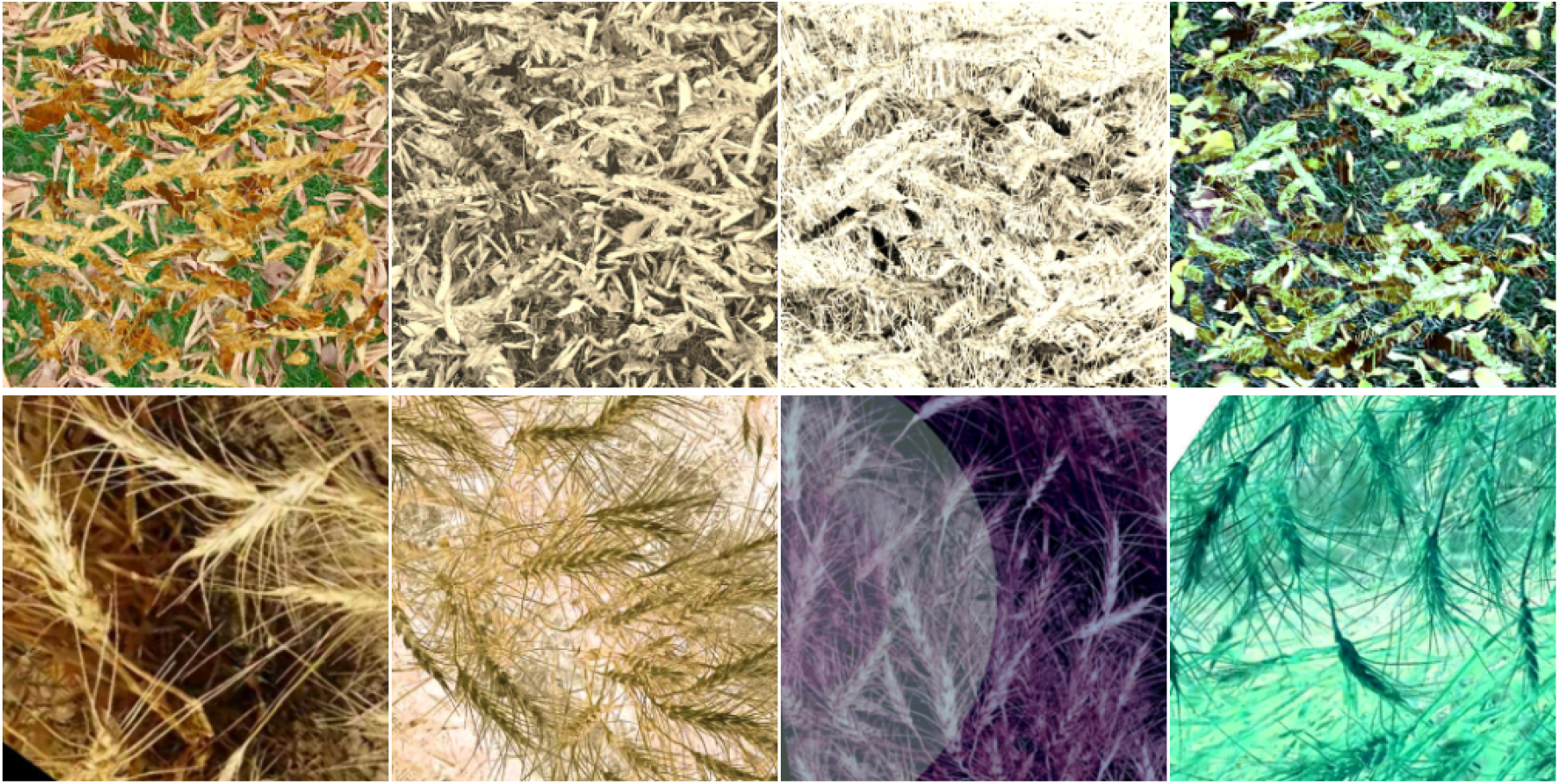
The top row illustrates examples of simulated images, and the bottom row shows examples of strongly augmented images generated in the first domain adaptation step.

**Figure 7:**
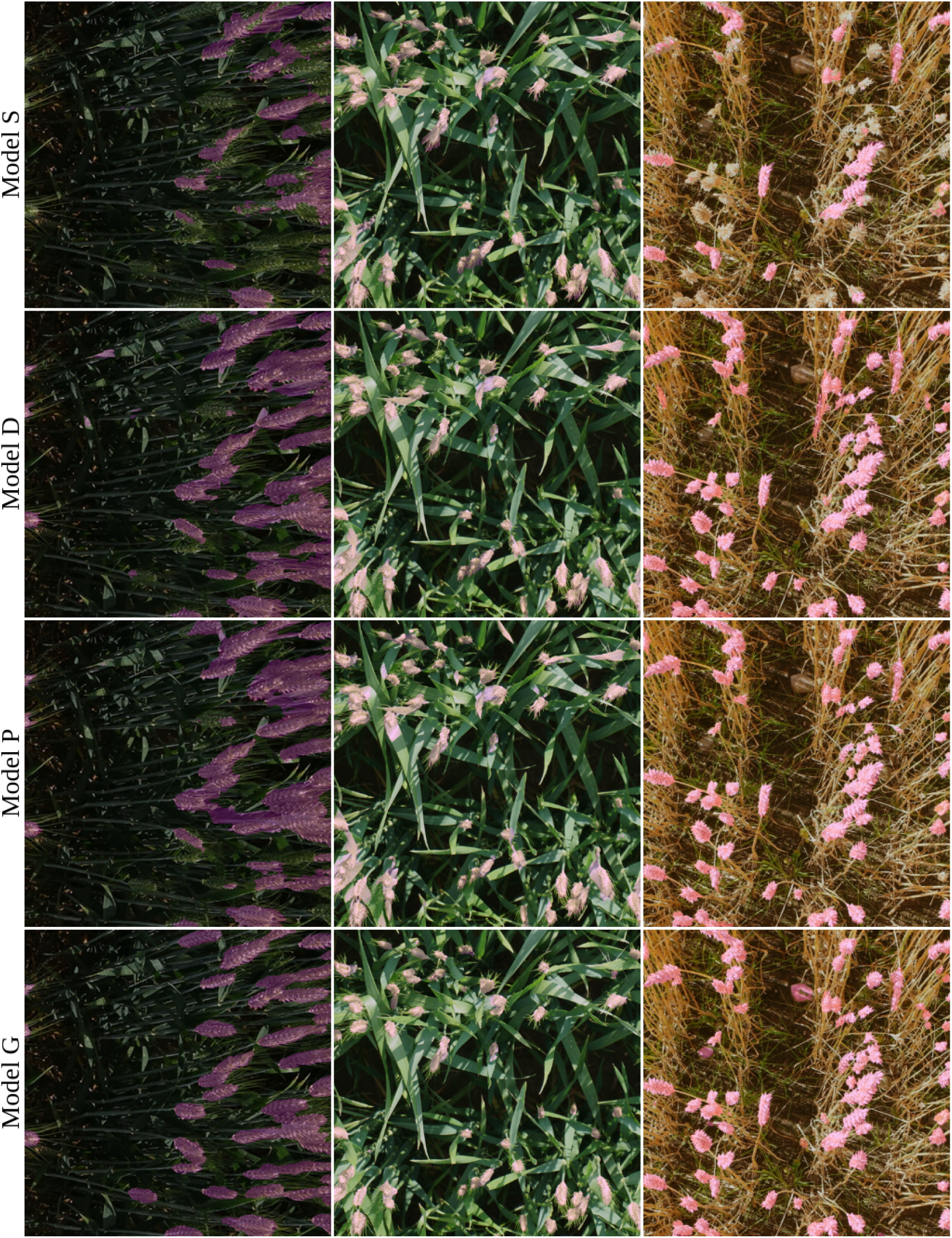
Examples of images from the GWHD dataset predicted by different models. The predicted masks are overlaid on the images. The images in each row shows the prediction made by a model. Model *S* is the model trained on the simulated dataset. Model *D, P*, and *G* are the results for the first, second, and third domain adaptation steps.

Even though datasets *D_t_* and *D_v_* better represent the real images of wheat fields, there is still a domain gap between the images in these datasets and real images. Therefore, we apply pseudo labeling as the second domain adaptation step. Using model *D*, we predict the segmentation mask for the images from the set *W*, excluding *w_t_, w_v_*, and all the test set image frames within 1 second from these images. We refer to these images as pseudo-labeled images since their masks might be noisy due to the model error. The resulting pseudo-labeled dataset is used to fine-tune model *D*. We refer to the model fine-tuned on the pseudo-labeled dataset as model *P*.

In the third domain adaptation step, we generate two sets *G_t_* and *G_v_*. We randomly select two images from each domain of the GWHD dataset, of which one is added to *G_t_* and the other is added to *G_v_*. Using model *P*, we predict a segmentation map for each image. If required, the predicted segmentation maps are then manually corrected. We also add *w_t_* to *G_t_* and *w_v_* to *G_v_*. We further expand *G_t_* by adding its 359 rotations of each image in *G_t_. G_v_* is also expanded analogously. These images are used to fine-tune model *P*. We refer to the fine-tuned model as model *G*.

To summarize the model development process, first, we use the datasets *S_t_* (for training) and *S_v_* (for validation) to develop model *S*. This model is then fine-tuned using datasets *D_t_* (for training) and *D_v_* (for validation) to develop model *D*. Next, we fine-tune model D using datasets *P_t_* (for training) and *P_v_* (for validation) to develop model *P*. Finally, we fine-tune model *P* using datasets *G_t_* (for training) and *G_v_* (for validation) to develop model *G* as the final model.

## 3 Results

Table 1 highlights the performance measures for the models on our internal test set, i.e., randomly chosen images from the wheat field video clip’s frames. The model developed utilizing two annotated images from a wheat field and short unannotated video clips of the wheat field and background fields achieved a high Dice score of 0.89 for segmenting the images from the same wheat field.

**Table 1:** The IoU and Dice score for the model *S*, which is trained on the simulated datasets, as well as models *D, P*, and *G* which are the models resulting from the first, second, and third domain adaptation steps. All models were evaluated using the internal test set, i.e., images extracted from the video clip of the wheat field. These performance measures are calculated for these models with and without test time augmentation (TTA). ImageNet refers to the initial model pretrained on the ImageNet dataset.

| Experiment | Initial Model | Dice | IoU | TTA Dice | TTA IoU |
| --- | --- | --- | --- | --- | --- |
| model $S$ | ImageNet | 0.7090 | 0.5658 | 0.7804 | 0.6463 |
| model $D$ | model $S$ | 0.8860 | 0.7974 | 0.8822 | 0.7910 |
| model $P$ | model $D$ | 0.8984 | 0.8167 | 0.8864 | 0.7975 |
| model $G$ | model $P$ | 0.8902 | 0.8038 | 0.8871 | 0.7986 |

When utilizing such a model as a general-purpose model for segmenting images in the GWHD dataset, the Dice score dropped to 0.73. This highlights the need to address the substantial domain shift caused by high variations in factors such as wheat growth stages, illumination, and imaging hardware and techniques. Table 2 shows the performance measures for the models evaluated using the GWHD dataset.

**Table 2:** The IoU and Dice score for the model *S*, which is trained on the simulated datasets, as well as models *D*, *P*, and *G*, which are the models resulting from the first, second, and third domain adaptation steps. All models were evaluated using the images from the GWHD dataset. These images have not been used for model development. The performance measures were calculated for these models with and without test time augmentation (TTA). ImageNet refers to the initial model pretrained on the ImageNet dataset.

| Experiment | Initial Model | Dice | IoU | TTA Dice | TTA IoU |
| --- | --- | --- | --- | --- | --- |
| model $S$ | ImageNet | 0.3678 | 0.2742 | 0.4845 | 0.3745 |
| model $D$ | model $S$ | 0.6528 | 0.5272 | 0.7416 | 0.6225 |
| model $P$ | model $D$ | 0.6748 | 0.5606 | 0.7289 | 0.6165 |
| model $G$ | model $P$ | 0.9144 | 0.8583 | 0.9084 | 0.8489 |

The GWHD dataset includes images from 18 different domains from various growth stages and imaging characteristics. Tables 3 and 4 show the performance measures for the trained models after applying the test time augmentation on each domain separately. As shown, the model performance varies across domains.

**Table 3:**
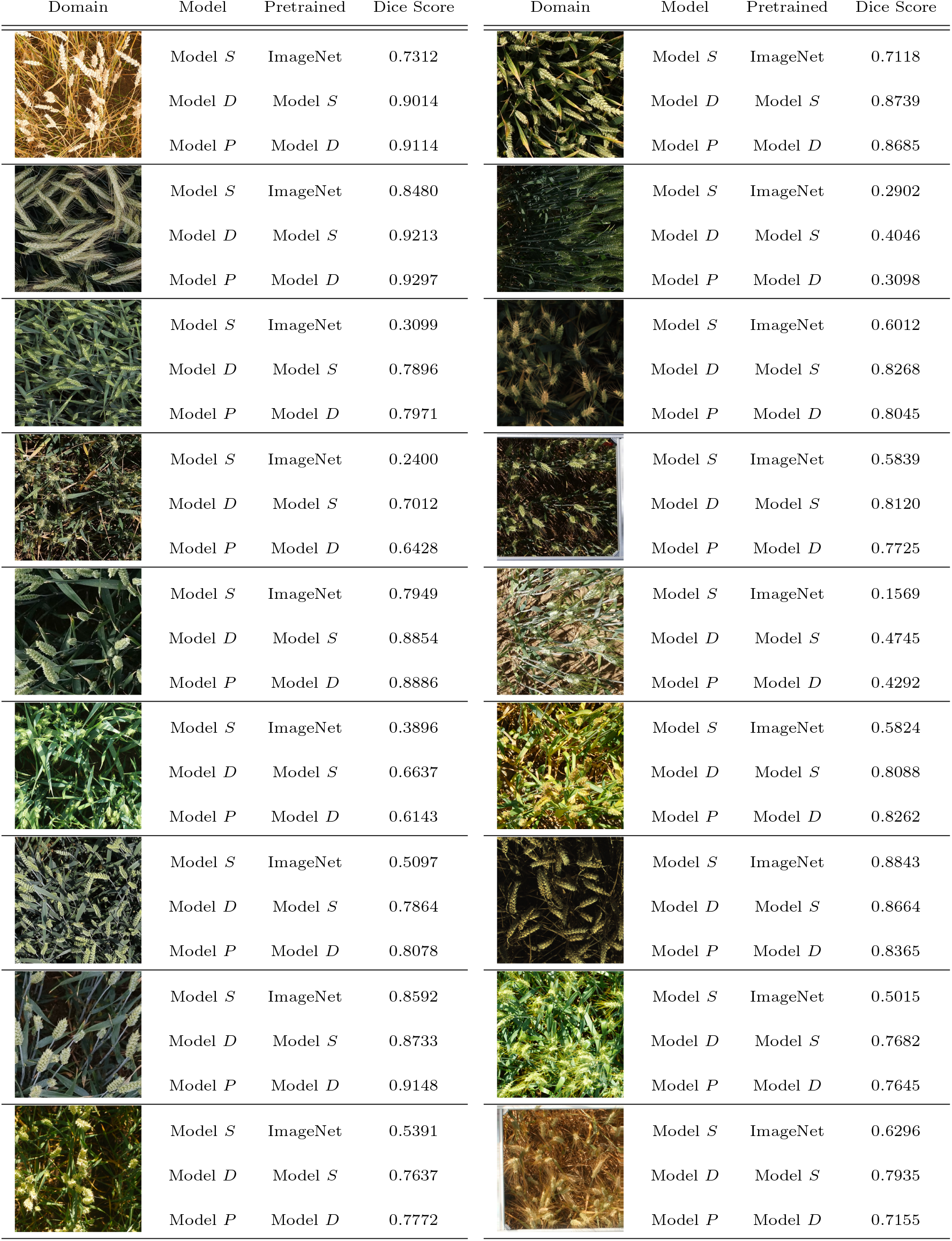
The performance of the models after applying the test time augmentation on each of the 18 domains of the GWHD dataset. Model *S* is the model trained using the simulated dataset. Models *D* and *P* are the models resulting from the first and second domain adaptation steps.

| Domain | Model | Pretrained | Dice Score | Domain | Model | Pretrained | Dice Score |
| --- | --- | --- | --- | --- | --- | --- | --- |
| | Model $S$ | ImageNet | 0.7312 | | Model $S$ | ImageNet | 0.7118 |
| | Model $D$ | Model $S$ | 0.9014 | | Model $D$ | Model $S$ | 0.8739 |
| | Model $P$ | Model $D$ | 0.9114 | | Model $P$ | Model $D$ | 0.8685 |
| | Model $S$ | ImageNet | 0.8480 | | Model $S$ | ImageNet | 0.2902 |
| | Model $D$ | Model $S$ | 0.9213 | | Model $D$ | Model $S$ | 0.4046 |
| | Model $P$ | Model $D$ | 0.9297 | | Model $P$ | Model $D$ | 0.3098 |
| | Model $S$ | ImageNet | 0.3099 | | Model $S$ | ImageNet | 0.6012 |
| | Model $D$ | Model $S$ | 0.7896 | | Model $D$ | Model $S$ | 0.8268 |
| | Model $P$ | Model $D$ | 0.7971 | | Model $P$ | Model $D$ | 0.8045 |
| | Model $S$ | ImageNet | 0.2400 | | Model $S$ | ImageNet | 0.5839 |
| | Model $D$ | Model $S$ | 0.7012 | | Model $D$ | Model $S$ | 0.8120 |
| | Model $P$ | Model $D$ | 0.6428 | | Model $P$ | Model $D$ | 0.7725 |
| | Model $S$ | ImageNet | 0.7949 | | Model $S$ | ImageNet | 0.1569 |
| | Model $D$ | Model $S$ | 0.8854 | | Model $D$ | Model $S$ | 0.4745 |
| | Model $P$ | Model $D$ | 0.8886 | | Model $P$ | Model $D$ | 0.4292 |
| | Model $S$ | ImageNet | 0.3896 | | Model $S$ | ImageNet | 0.5824 |
| | Model $D$ | Model $S$ | 0.6637 | | Model $D$ | Model $S$ | 0.8088 |
| | Model $P$ | Model $D$ | 0.6143 | | Model $P$ | Model $D$ | 0.8262 |
| | Model $S$ | ImageNet | 0.5097 | | Model $S$ | ImageNet | 0.8843 |
| | Model $D$ | Model $S$ | 0.7864 | | Model $D$ | Model $S$ | 0.8664 |
| | Model $P$ | Model $D$ | 0.8078 | | Model $P$ | Model $D$ | 0.8365 |
| | Model $S$ | ImageNet | 0.8592 | | Model $S$ | ImageNet | 0.5015 |
| | Model $D$ | Model $S$ | 0.8733 | | Model $D$ | Model $S$ | 0.7682 |
| | Model $P$ | Model $D$ | 0.9148 | | Model $P$ | Model $D$ | 0.7645 |
| | Model $S$ | ImageNet | 0.5391 | | Model $S$ | ImageNet | 0.6296 |
| | Model $D$ | Model $S$ | 0.7637 | | Model $D$ | Model $S$ | 0.7935 |
| | Model $P$ | Model $D$ | 0.7772 | | Model $P$ | Model $D$ | 0.7155 |

**Table 4:**
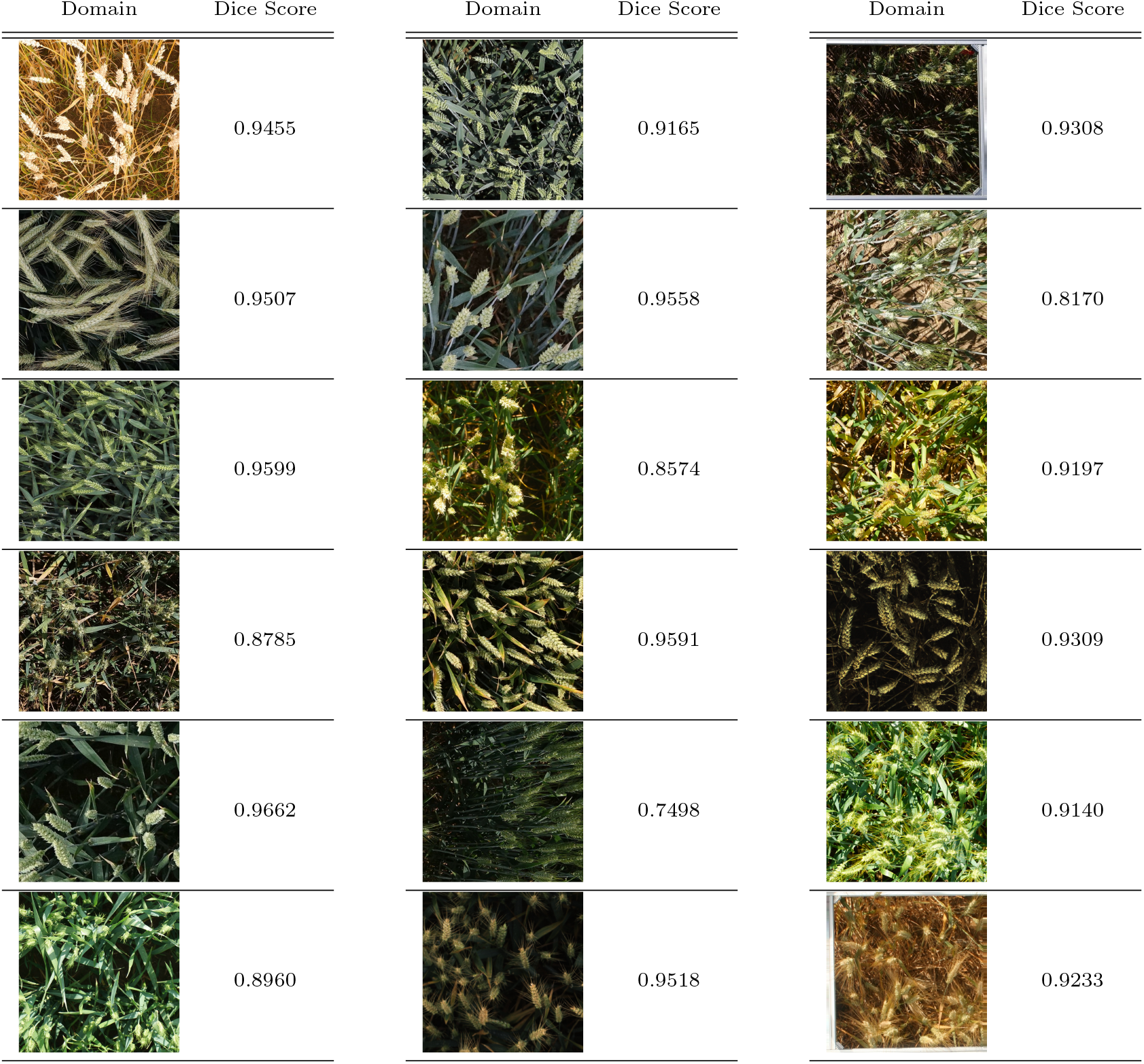
The performance of the models after applying the test time augmentation on each of the 18 domains of the GWHD dataset. Model *G* is the resulting model after the third domain adaptation step, i.e., the model pretrained on model *P*, and fine-tuned on the 18 images selected from GWHD (one per domain).

| Domain | Dice Score | Domain | Dice Score | Domain | Dice Score |
| --- | --- | --- | --- | --- | --- |
|  | 0.9455 |  | 0.9165 |  | 0.9308 |
|  | 0.9507 |  | 0.9558 |  | 0.8170 |
|  | 0.9599 |  | 0.8574 |  | 0.9197 |
|  | 0.8785 |  | 0.9591 |  | 0.9309 |
|  | 0.9662 |  | 0.7498 |  | 0.9140 |
|  | 0.8960 |  | 0.9518 |  | 0.9233 |

## 4 Discussion

In this study, we proposed a semi-self-supervised learning approach to tackle the wheat head semantic segmentation problem. Our approach, based on simulating computationally-annotated datasets followed by domain adaptation steps, makes it possible to develop a high-performing semantic segmentation model for wheat head segmentation using only two manually annotated images per domain. This allows for the adoption of deep learning technology in similar applications where a large-scale annotated dataset is unavailable due to the costly, tedious, and time-consuming nature of manual annotation. Our final model achieved the state-of-the-art performance for wheat head segmentation as it reached a Dice score of 0.91 on the GWHD dataset, which is a diverse set of images from five countries and 18 different domains. Also, the proposed approach is model agnostic and can be further improved by using different deep learning model architectures.

The model trained only using the simulated datasets achieved a dice score of 0.78 on the internal single domain dataset. However, its performance dropped to 0.48 when evaluated on our external and diverse test set. This underscores the effect of domain shift between simulated and real images. The substantial improvement in the performance measures after applying the first and second domain adaptation steps further highlights the utility of these domain adaptation steps in filling the gap between simulated and real images.

Even after applying the second domain adaptation step, we observed a drop in performance measures when we applied the resulting model to our external test set. This gap can be attributed to the domain shift between the images from the single wheat field used for model training and images from other domains, e.g., different growth stages or environmental conditions. This suggests that models trained on data from a single domain might not generalize to external data. Further, internal evaluation using a test set that comes from the same distribution as the training set often does not provide a reliable estimate of generalization error. Using multi-domain and diverse datasets for model evaluation should be considered the best practice.

In this study, we only utilized a short video clip of a wheat field to develop a deep semantic segmentation model. However, utilizing a larger number of video clips of wheat fields from various growth stages of the wheat could potentially further improve the performance of the model trained only using computationally-annotated datasets.

In addition to the single wheat field video, we used a group of 11 background scene videos in which there is no frame capturing wheat heads. In the data simulation step, we overlaid wheat heads onto image frames extracted from these background videos. By utilizing various background videos, we primarily intended to increase data variation reducing the chance of overfitting. We suggest a comprehensive study of the effect of various background videos on the model performance as future research.

We showed that the proposed approach could lead to developing high-performing models for semantic segmentation of wheat heads. However, the application of the proposed method is not limited to the wheat head segmentation, and it could be used for other applications such as segmenting other crops where the aim is to segment a highly self-similar patterns such as leaves, spikes, flowers, or fruits. Note that while a single image could provide a good representation of a wheat field, it does not offer a good representation of typical pictures such as those in the ImageNet dataset. Therefore, one should not expect to use one sample from the ImageNet dataset to provide a segmentation model for images in ImageNet.

In this paper, we only used a single video clip of wheat field in the final growth stage. This led to a high-performing model for segmenting images from the same field and the same growth stage. However, when the model was applied to the Global Wheat Head Detection dataset—a highly diverse dataset of wheat field images from five countries and 18 domains representing various growth stages and imaging conditions—the model performance slightly decreased. This highlights that developing a general model that can optimally work for different imaging conditions and growth stages requires images representing such conditions. The proposed approach could facilitate creating such a dataset by utilizing video clips instead of large datasets of images taken individually and then only annotating a single image from each video clip. We suggest using a set of video clips representing various growth stages of the wheat for future research.

Also, in this paper, we used a modified U-Net model architecture for all experiments. However, the proposed approach is independent of the model architecture and can benefit from different model architectures. We suggest utilizing different model architectures for future research.

## 5 Conclusions

In this work, we introduced a semi-self-supervised approach for the wheat head semantic segmentation task by simulating a computationally-annotated large-scale dataset and applying three domain adaptation steps for addressing domain shift. Utilizing a few short video clips of a wheat field and background vegetation, the proposed method facilitates data collection for model building. Further, since the proposed model only uses a few manually annotated images, it avoids the manual annotation of a large dataset, facilitating model development. While we showed the utility of the proposed method for wheat head segmentation, it could be applied to other applications that have similar dense repeating patterns of objects, such as segmenting plant organ is other crop species, or segmenting molecular components in microscopy images.

## Appendix A

In this study, we benefit from a variety of image augmentations, including pixel-level and spatial-level transformations from the Albumentations package [51]. To simulate *S_t_* and *S_v_*, real and fake wheat heads undergo a sequence of transformations including HorizontalFlip, VerticalFlip, Rotate, and ElasticTransform. After overlapping both real and fake wheat heads on the background images, we augment the resulting images using a long list of pixel-level transformations such as ColorJitter, ChannelShuffle, RGBShift, ChannelDropout, HueSaturationValue, Emboss, Solarize, InvertImg, ToGray, ToSepia, FancyPCA, Posterize, Sharpen, RandomGamma, Equalize, RandomBrightnessContrast, CLAHE, GaussianBlur, MotionBlur, RandomRain, RandomFog, RandomSnow, RandomSunFlare, GaussNoise, MultiplicativeNoise, ISONoise, and Normalize.

When generating *D_t_, D_v_*, in addition to the color transformations applied to the simulated dataset, we included the following image augmentations: Flip, Rotation, ElasticTransform, GridDistortion, as well as a long list of RandomCrop functions with different squared crop sizes ranging from 400 × 400 to 1000 × 1000. These images then were resized to 1024 × 1024.

In the pseudo-labeling step, we tried all of the augmentation methods used when generating *D_t_* and *D_v_* (i.e., the first domain adaptation step) but excluded those that drastically changed the color, like ColorJitter, Channel Shuffle, and RGB Shift. In the last training step, we also utilized the same list applied in the first domain adaptation step to augment the chosen training samples from the GWHD dataset. During model evaluation, only the Resize, and Normalize transformations were applied to images.

## References

[1] W. Wang, Y. Yang, X. Wang, W. Wang, and J. Li, “Development of convolutional neural network and its application in image classification: A survey,” Optical Engineering, vol. 58, no. 4, p. 040901, 2019.

[2] L. Liu et al., “Deep learning for generic object detection: A survey,” International Journal of Computer Vision, vol. 128, no. 2, pp. 261–318, 2020.

[3] A. M. Hafiz and G. M. Bhat, “A survey on instance segmentation: State of the art,” International Journal of Multimedia Information Retrieval, vol. 9, no. 3, pp. 171–189, 2020.

[4] S. Hao, Y. Zhou, and Y. Guo, “A brief survey on semantic segmentation with deep learning,” Neurocomputing, vol. 406, pp. 302–321, 2020.

[5] J. R. Ubbens and I. Stavness, “Deep plant phenomics: A deep learning platform for complex plant phenotyping tasks,” Frontiers in Plant Science, vol. 8, p. 1190, 2017.

[6] Y.-Y. Zheng, J.-L. Kong, X.-B. Jin, X.-Y. Wang, T.-L. Su, and M. Zuo, “CropDeep: The crop vision dataset for deep-learning-based classification and detection in precision agriculture,” Sensors, vol. 19, no. 5, p. 1058, 2019.

[7] X.-B. Jin, X.-H. Yu, X.-Y. Wang, Y.-T. Bai, T.-L. Su, and J.-L. Kong, “Deep learning predictor for sustainable precision agriculture based on internet of things system,” Sustainability, vol. 12, no. 4, p. 1433, 2020.

[8] K. He, X. Zhang, S. Ren, and J. Sun, “Deep residual learning for image recognition,” in Proceedings of the IEEE Conference on Computer Vision and Pattern Recognition, 2016, pp. 770–778.

[9] S. Ren, K. He, R. Girshick, and J. Sun, “Faster R-CNN: Towards real-time object detection with region proposal networks,” Advances in Neural Information Processing Systems, vol. 28, pp. 91–99, 2015.

[10] K. He, G. Gkioxari, P. Dollár, and R. Girshick, “Mask R-CNN,” in Proceedings of the IEEE International Conference on Computer Vision, 2017, pp. 2961–2969.

[11] S. Bhagat, M. Kokare, V. Haswani, P. Hambarde, and R. Kamble, “WheatNet-Lite: A novel light weight network for wheat head detection,” in Proceedings of the IEEE/CVF International Conference on Computer Vision, 2021, pp. 1332–1341.

[12] S. Mardanisamani and M. Eramian, “Segmentation of vegetation and microplots in aerial agriculture images: A survey,” The Plant Phenome Journal, vol. 5, no. 1, e20042, 2022.

[13] E. David et al., “Global wheat head detection 2021: An improved dataset for benchmarking wheat head detection methods,” Plant Phenomics, vol. 2021, 2021.

[14] S. Khaki, N. Safaei, H. Pham, and L. Wang, “WheatNet: A lightweight convolutional neural network for high-throughput image-based wheat head detection and counting,” Neurocomputing, vol. 489, pp. 78–89, 2022.

[15] F. Han and J. Li, “Wheat heads detection via yolov5 with weighted coordinate attention,” in 2022 7th International Conference on Cloud Computing and Big Data Analytics (ICCCBDA), IEEE, 2022, pp. 300–306.

[16] B. Gong, D. Ergu, Y. Cai, and B. Ma, “Real-time detection for wheat head applying deep neural network,” Sensors, vol. 21, no. 1, p. 191, 2020.

[17] A. Bochkovskiy, C.-Y. Wang, and H.-Y.M. Liao, “YOLOv4: Optimal speed and accuracy of object detection,” arXiv preprint arXiv:2004.10934, 2020.

[18] J. Redmon and A. Farhadi, “YOLOv3: An incremental improvement,” arXiv preprint arXiv: 1804.02767, 2018.

[19] K. He, X. Zhang, S. Ren, and J. Sun, “Spatial pyramid pooling in deep convolutional networks for visual recognition,” IEEE Transactions on Pattern Analysis and Machine Intelligence, vol. 37, no. 9, pp. 1904–1916, 2015.

[20] C.-Y. Wang, H.-Y.M. Liao, Y.-H. Wu, P.-Y. Chen, J.-W. Hsieh, and I.-H. Yeh, “Cspnet: A new backbone that can enhance learning capability of cnn,” in Proceedings of the IEEE/CVF Conference on Computer Vision and Pattern Recognition Workshops, 2020, pp. 390–391.

[21] C. Liu, K. Wang, H. Lu, and Z. Cao, “Dynamic color transform for wheat head detection,” in Proceedings of the IEEE/CVF International Conference on Computer Vision, 2021, pp. 1278–1283.

[22] M. Sandler, A. Howard, M. Zhu, A. Zhmoginov, and L.-C. Chen, “MobileNetV2: Inverted residuals and linear bottlenecks,” in Proceedings of the IEEE Conference on Computer Vision and Pattern Recognition, 2018, pp. 4510–4520.

[23] S. Rawat, A.L. Chandra, S.V. Desai, V.N. Balasubramanian, S. Ninomiya, and W. Guo, “How useful is image-based active learning for plant organ segmentation?” Plant Phenomics, vol. 2022, 2022.

[24] B.R. Hussein, O.A. Malik, W.-H. Ong, and J.W.F. Slik, “Automated extraction of phenotypic leaf traits of individual intact herbarium leaves from herbarium specimen images using deep learning based semantic segmentation,” Sensors, vol. 21, no. 13, p. 4549, 2021.

[25] L.-C. Chen, Y. Zhu, G. Papandreou, F. Schroff, and H. Adam, “Encoder-decoder with atrous separable convolution for semantic image segmentation,” in Proceedings of the European Conference on Computer Vision (ECCV), 2018, pp. 801–818.

[26] K. Simonyan and A. Zisserman, “Very deep convolutional networks for large-scale image recognition,” arXiv preprint arXiv:1409.1556, 2014.

[27] T. Alkhudaydi, D. Reynolds, S. Griffiths, J. Zhou, and B. De La Iglesia, “An exploration of deep-learning based phenotypic analysis to detect spike regions in field conditions for UK bread wheat,” Plant Phenomics, vol. 2019, 2019.

[28] J. Long, E. Shelhamer, and T. Darrell, “Fully convolutional networks for semantic segmentation,” in Proceedings of the IEEE Conference on Computer Vision and Pattern Recognition, 2015, pp. 3431–3440.

[29] P. Sadeghi-Tehran, N. Virlet, E.M. Ampe, P. Reyns, and M.J. Hawkesford, “DeepCount: In-field automatic quantification of wheat spikes using simple linear iterative clustering and deep convolutional neural networks,” Frontiers in Plant Science, vol. 10, p. 1176, 2019.

[30] J. Ma et al., “Improving segmentation accuracy for ears of winter wheat at flowering stage by semantic segmentation,” Computers and Electronics in Agriculture, vol. 176, p. 105662, 2020.

[31] X. Xiong et al., “Panicle-SEG: A robust image segmentation method for rice panicles in the field based on deep learning and superpixel optimization,” Plant Methods, vol. 13, no. 1, pp. 1–15, 2017.

[32] C. Tan et al., “Rapid recognition of field-grown wheat spikes based on a superpixel segmentation algorithm using digital images,” Frontiers in Plant Science, vol. 11, p. 259, 2020.

[33] R. Achanta, A. Shaji, K. Smith, A. Lucchi, P. Fua, and S. Süsstrunk, “Slic superpixels compared to state-of-the-art superpixel methods,” IEEE Transactions on Pattern Analysis and Machine Intelligence, vol. 34, no. 11, pp. 2274–2282, 2012.

[34] L. Schmarje, M. Santarossa, S.-M. Schröder, and R. Koch, “A survey on semi-, self-and unsupervised learning for image classification,” IEEE Access, vol. 9, pp. 82146–82168, 2021.

[35] X.J. Zhu, “Semi-supervised learning literature survey,” 2005.

[36] Y.-H.H. Tsai, Y. Wu, R. Salakhutdinov, and L.-P. Morency, “Self-supervised learning from a multi-view perspective,” arXiv preprint arXiv:2006.05576, 2020.

[37] N. Komodakis and S. Gidaris, “Unsupervised representation learning by predicting image rotations,” in International Conference on Learning Representations (ICLR), 2018.

[38] D. Pathak, P. Krahenbuhl, J. Donahue, T. Darrell, and A. A. Efros, “Context encoders: Feature learning by inpainting,” in Proceedings of the IEEE Conference on Computer Vision and Pattern Recognition, 2016, pp. 2536–2544.

[39] M. Noroozi and P. Favaro, “Unsupervised Learning of Visual Representations by Solving Jigsaw Puzzles,” in European Conference on Computer Vision, Springer, 2016, pp. 69–84.

[40] C. Wei et al., “Iterative reorganization with weak spatial constraints: Solving arbitrary jigsaw puzzles for unsupervised representation learning,” in Proceedings of the IEEE/CVF Conference on Computer Vision and Pattern Recognition, 2019, pp. 1910–1919.

[41] D. Kim, D. Cho, D. Yoo, and I.S. Kweon, “Learning image representations by completing damaged jigsaw puzzles,” in 2018 IEEE Winter Conference on Applications of Computer Vision (WACV), IEEE, 2018, pp. 793–802.

[42] F. Fourati, W.S. Mseddi, and R. Attia, “Wheat head detection using deep, semi-supervised and ensemble learning,” Canadian Journal of Remote Sensing, vol. 47, no. 2, pp. 198–208, 2021.

[43] M. Tan, R. Pang, and Q. V. Le, “Efficientdet: Scalable and efficient object detection,” in Proceedings of the IEEE/CVF Conference on Computer Vision and Pattern Recognition, 2020, pp. 10 781–10 790.

[44] K. Najafian, A. Ghanbari, I. Stavness, L. Jin, G. H. Shirdel, and F. Maleki, “A Semi-Self-Supervised learning approach for wheat head detection using extremely small number of labeled samples,” in Proceedings of the IEEE/CVF International Conference on Computer Vision,2021, pp. 1342–1351.

[45] M. Everingham, L. Van Gool, C.K. Williams, J. Winn, and A. Zisserman, “The pascal visual object classes (voc) challenge,” International Journal of Computer Vision, vol. 88, no. 2, pp. 303–338, 2010.

[46] T.-Y. Lin et al., “Microsoft coco: Common objects in context,” in European Conference on Computer Vision, Springer, 2014, pp. 740–755.

[47] O. Ronneberger, P. Fischer, and T. Brox, “U-Net: Convolutional networks for biomedical image segmentation,” in International Conference on Medical Image Computing and Computer-assisted Intervention, Springer, 2015, pp. 234–241.

[48] M. Tan and Q. Le, “EfficientNet: Rethinking model scaling for convolutional neural networks,” in International Conference on Machine Learning, PMLR, 2019, pp. 6105–6114.

[49] A. Krizhevsky, I. Sutskever, and G.E. Hinton, “Imagenet Classification with Deep Convolutional Neural Networks,” Advances in Neural Information Processing Systems, vol. 25, pp. 1097–1105, 2012.

[50] P. Yakubovskiy, Segmentation models pytorch, https://github.com/qubvel/segmentation_models.pytorch, 2020.

[51] A. Buslaev, V.I. Iglovikov, E. Khvedchenya, A. Parinov, M. Druzhinin, and A.A. Kalinin, “Albumentations: Fast and flexible image augmentations,” Information, vol. 11, no. 2, 2020, issn:2078-2489. doi:10.3390/info11020125. [Online]. Available: https://www.mdpi.com/2078-2489/11/2/125.

[52] J. Bertels et al., “Optimizing the dice score and jaccard index for medical image segmentation: Theory and practice,” in International Conference on Medical Image Computing and Computer-assisted Intervention, Springer, 2019, pp. 92–100.

[53] I. Joshi et al., “Explainable fingerprint roi segmentation using monte carlo dropout,” in Proceedings of the IEEE/CVF Winter Conference on Applications of Computer Vision, 2021, pp. 60–69.

[54] S. Ruder, “An overview of gradient descent optimization algorithms,” arXiv preprint arXiv: 1609.04747, 2016.

[55] P. Simard, D. Steinkraus, and J. Platt, “Best practices for convolutional neural networks applied to visual document analysis,” in Seventh International Conference on Document Analysis and Recognition, 2003. Proceedings., vol. 3, 2003, pp. 958–963. doi:10.1109/ICDAR.2003.1227801.

